# Room Temperature Isothermal Colorimetric Padlock Probe Rolling Circle Amplification for Viral DNA and RNA Detection

**DOI:** 10.1101/2020.06.12.128876

**Authors:** Wilson Huang, Joyce Ting, Matthew Fang, Hannah Hsu, Jimmy Su, Tsuyoshi Misaki, Derek Chan, Justin Yang, Ting-Yu Yeh, Kelly Yang, Vera Chien, Tiffany Huang, Andrew Chen, Claire Wei, Jonathan Hsu, Jude C. Clapper

## Abstract

Seasonal flu and pandemics, which account for millions of infections and hundreds of thousands of deaths, require rapid and reliable detection mechanisms to implement preventive and therapeutic measures. Current detection methods of viral infections have limitations in speed, accuracy, accessibility, and usability. This project presents a novel, widely applicable viral diagnostic test that uses a modified version of rolling circle amplification (RCA) to be sensitive, specific, direct RNA targeted, colorimetric and operable at room temperature. We are specifically detecting the following high-impact viruses: SARS-CoV-2, Influenza A (H1N1pdm09), and Influenza B (Victoria Lineage), although our test can be adapted to any viral infection. Results using synthetic viral DNA and RNA sequences show that our diagnostic test takes approximately one hour, detects femtomolar concentrations of RNA strands, and differentiates between virus strains. We believe implementing our diagnostic test will provide faster responses to future viral-related outbreaks for quicker societal recovery.

## Introduction

### Virus Related Infections and Deaths

Seasonal flu accounts for approximately 3 to 5 million cases and 250 to 500 thousands deaths worldwide annually (*Influenza (Seasonal)*, n.d.). Pandemics such as the ongoing SARS-CoV2 pandemic have already caused 108.19 million cases and 2.38 million deaths, and the numbers continue to rise (*Microsoft Bing COVID-19 Tracker*, n.d.). The number of people subjected to these diseases makes diagnostic measures crucial to installing preventive and therapeutic action. Our study aims to detect SARS-CoV-2, Influenza A (H1N1pdm09), and Influenza B (Victoria Lineage), due to their high impact factor in both infections and deaths.

### Current Methods of Sample Collection and Purification

In order to test for a viral infection, a sample must be collected and at times isolated. Currently, there are two main methods for viral collection: through nasopharyngeal swabs or saliva. For nasopharyngeal swabs, a swab is inserted into the nostril until it reaches the pharynx region. It is left there for a few seconds to absorb secretions and removed while slowly rotating to obtain cells (Marty et al., 2020). This is usually done by a medical personnel. Saliva on the other hand is usually self collected by spitting into a sterile container with saline (Jamal et al., n.d.). Various articles have shown that saliva collection is highly sensitive and yields similar test outcomes, making it a reliable, noninvasive alternative to swabs (Greenwood, 2020; Iwasaki et al., 2020; Jamal et al., n.d.; *Nasopharyngeal Swab vs. Saliva for COVID-19 Diagnosis*, 2020).

Most viral extraction or purification techniques center around commercial RNA extraction kits. They usually involve using lysis buffers, wash buffers, elution buffers, membrane columns, and centrifugation to ultimately extract in a lengthy process (Tan & Yiap, 2009). These are more often for nucleic acid tests and aren’t usually required in rapid test settings.

### Reverse Transcriptase Polymerase Chain Reaction

The gold standard of viral detection is reverse transcriptase polymerase chain reaction (RT-PCR)(CDC, 2020). This methodology allows for high accuracy and sensitivity, but due to the need of varying temperatures and thus a thermocycler, this option is not only expensive but also requires skilled technicians to run tests and interpret the data (Corman et al., 2020; da Costa Lima et al., 2013; *False-Negative Rate of RT-PCR SARS-CoV-2 Tests*, n.d.; Tahamtan & Ardebili, 2020). RT-PCR also has limited specificity due to the reliance of reverse transcriptase, which has no proofreading capacity, to form cDNA sequences (Alhassan et al., 2015).

Approximately 1 error is inputted for every 1700 polymerized nucleotides but can reach as high as 1 error per 70 polymerized nucleotides. RT-PCR compared to other methods of detection is relatively accurate (Bhagavan & Ha, 2015; Roberts et al., 1988). However, even the lowest rate of transcription error through reverse-transcriptase is 21% after 8 days of virus exposure and can go as high as 67% within the first 5 days after virus exposure (*False-Negative Rate of RT-PCR SARS-CoV-2 Tests*, n.d.).

### Isothermal Amplification - Loop Mediated Isothermal Amplification

A rising method of nucleic acid amplification is isothermal amplification, which allows the amplification reaction to run at a constant temperature. One isothermal technique is loop mediated isothermal amplification (LAMP). LAMP utilizes 4-6 sets of primers to amplify many regions of the target sequence to ultimately create a loop structure that can be used as a backbone for further exponential amplification (Biolabs, n.d.). Real time fluorescent detection through probes, lateral flow, or DNA electrophoresis can then be used to examine the amplified or unamplified product (Biolabs, n.d.). However, one issue with LAMP is its need to run at 65°C (Ge et al., 2017). This requires the use of an incubator or thermocycler, which defeats the purpose of using LAMP instead of RT-PCR since specific instrumentation or even the same machinery not so accessible may have to be used. LAMP also relies on reverse transcriptase for RNA virus detection, which limits the specificity of the assay due to the low fidelity of reverse transcriptase (Roberts et al., 1988).

### Protein/Antibody Tests - Lateral Flow Immunoassay

A form of rapid test is through lateral flow immunoassays. On the top of the strip is usually an antibody whereas the bottom of the strip usually contains gold nanoparticles with either antibodies or viral antigens expressed on it (Koczula & Gallotta, 2016; Sajid et al., 2015). If viral antigens are being detected, the antibody nanoparticle that captures the antigen carries it to the detecting antibody (Koczula & Gallotta, 2016; Sajid et al., 2015). If human antibodies are being detected, then the viral antigen nanoparticle will capture the antibody and the detecting antibody will capture the nanoparticle antigens (Koczula & Gallotta, 2016; Sajid et al., 2015).

Whatever the case, the presence of virus particles with the expressed antigens or human antibodies in response to virus exposure will cause specific lines to appear as a result of captured gold nanoparticles along the lines of capture antibodies (Koczula & Gallotta, 2016). These rapid tests do not require sample preparation and provide fast results. However, due to the lack of signal amplification of any kind, it’s usually low in sensitivity and highly inaccurate (Gordon & Michel, 2008; *Rapid Influenza Diagnostic Tests* | *CDC*, 2019). Antibody testing often leads to a delayed diagnosis, as it takes approximately 1 to 2 weeks for humans to mount an immune response after the first onset of symptoms (Lippi et al., 2020).

### Protein/Antibody Tests - Blood Serology

Blood serology tests usually use the bioanalytical technique indirect ELISA. Plating the ELISA wells with viral antigens, a sample with human antibodies mounted in response to the virus will bind to the viral antigen (Dhamad & Abdal Rhida, 2020). A secondary antibody typically conjugated with an enzyme or fluorophore can then bind onto the Fc domains of the antibody sample to provide a colorimetric or fluorescent readout (Dhamad & Abdal Rhida, 2020). These ELISA plates can be scanned with a reader to quantify these values to be interpreted for results. However, the use of ELISA plates runs the risk of cross contamination and also requires a technician to follow a specific procedure to undertake the process (*ELISA Troubleshooting Tips* | *Abcam*, n.d.). As stated, there is also a risk of delayed diagnosis.

### Viral Cell Cultures

A method that is neither nucleic acid testing nor protein and antibody testing is viral cell culture (CDC, 2020). Through growth mediums such as RetroNectin, medical personnel attempt to culture viruses from isolated samples. In other cases, viruses are first used to transfection mammalian cell lines to allow for expansion before culture (Harcourt et al., n.d.). The growth of a virus doesn’t completely identify the type of virus a patient has however. To identify it, DNA sequencing, IF staining, electron microscopy, and other methods are employed (Harcourt et al., n.d.; Hematian et al., 2016). As one can expect, this process can take long, sometimes up to 30 days (Harcourt et al., n.d.). It also poses significant dangers to the individuals operating the test, if not handled correctly (*Virus Contaminations of Cell Cultures – A Biotechnological View*, n.d.).

### Our Selected Methodology

To address some of the issues with the aforementioned viral detection tests. We employed an isothermal methodology known as padlock probe rolling circle amplification. The two ends of the padlock probe are designed to bind to a specific complementary target sequence. Upon hybridization, a ligase, usually T4 ligase, can ligate the ends of the padlock probe due to a 5’ phosphorylation on the padlock probe and the presence of ATP as a cofactor (Hamidi & Ghourchian, 2015). The result is a circularized padlock probe, which can now act as a template for DNA amplification (Hamidi & Perreault, 2019). If the padlock probe is not complementary to the target sequence, ligation should not occur (Gu et al., 2018). Without a circularized padlock probe, continual DNA amplification will not be able to occur and thus the readout will show a negative result. The subsequent methods of amplification vary. Some involve the use of the 3’ to 5’ exonuclease activity of phi29 DNA polymerase to cut the target sequence until the ligation junction is reached, which enables DNA synthesis to proceed (Li et al., 2017). Others use an exonuclease or RNase to achieve the same function or direct annealing of a forward primer to the circularized backbone (Joffroy et al., 2018; *RNase H-Assisted RNA-Primed Rolling Circle Amplification for Targeted RNA Sequence Detection* | *Scientific Reports*, n.d.). Whatever the case, the DNA polymerase of choice, usually Bst DNA polymerase or phi29 DNA polymerase, has an existing DNA strand to synthesize DNA strands in circles around the padlock probe through its helicase activity (Joffroy et al., 2018). A second reverse primer that binds to the newly synthesized DNA strands is sometimes used to further increase DNA synthesis by introducing a branching-like amplification (Hamidi & Ghourchian, 2015; Hamidi & Perreault, 2019; Larsson et al., 2004). Finally, the readout usually uses fluorescent probes that recognize the synthesized strand and is thus quantifiable (Hamidi & Ghourchian, 2015; Larsson et al., 2004).

### Our Modifications to Traditional Rolling Circle Amplification

To maximize the Rolling Circle Amplification technique, we made modifications to increase accuracy through sensitivity and specificity and add additional functions that will increase usability. The modifications we included involve a colorimetric readout. We made use of the fact that for every nucleotide incorporated in DNA amplification, there is a release of H+ and PPi. Thus, with a pH indicator we can visualize a color change readout through a change in hydrogen ion concentration. But this also requires us to take out all buffers in the solutions of the reaction that might neutralize the hydrogen ions produced. Thus, our reaction was carried out with specially made solutions without any buffers such as Tris-HCl. Another modification is our use of splintR ligase (PBCV-1 DNA ligase and Chlorella virus DNA ligase) to allow the ligation of our DNA padlock probe and viral RNA target. This enables high specificity through direct RNA targeting. But to prevent RNA degradation, we used Protector Rnase Inhibitors to neutralize the activity of RNases that cleaves RNA. We also used phi29 DNA polymerase along with splintR ligase, as both enzymes operate at optimal settings near room temperature (30°C and 25°C respectively) and does in fact allow a room temperature reaction as experimentally proven. To enable the detachment of the target sequence to the circularized padlock probe in a simpler method, we designed universal primers that would bind to the backbone (non-target region) of the padlock probe. As a result, the helicase activity of the polymerase itself would remove the target sequence and allow for amplification to continue. An additional reverse primer that binds to the newly synthesized DNA strands is used to introduce a branching-like amplification. These methods ultimately help improve signal amplification which helps provide readouts even during low viral concentrations, increasing the sensitivity of our test. Studies have shown that the exonuclease activity of phi29 DNA polymerase can involve not only single stranded sequences but double stranded complementary sequences, we added phosphorothioate bonds to disable phi29 DNA polymerase from digesting it (Li et al., 2017).

All of these traits work together to form our simple, fast, and accurate universal viral detection platform.

## Methods

### Synthetic Viral Targets, Probes and Primers

For our padlock probe (PLP) design, we decided to target the Spike (S) glycoprotein gene for SARS-CoV-2 (C-19) and the Hemagglutinin (HA) gene for Influenza A (INFA) and B (INFB) due to their high copy number in viruses. In order to create a highly specific target sequence, the sequence had to be unique to each virus type while also being able to recognize subtle mutations over time. To that end, we aligned the nucleotide sequences of the S gene from several different SARS-CoV-2 variant strains as well as other related coronaviruses (such as SARS and MERS) across different dates. We identified a 36bp sequence that was highly conserved between various mutations within the SARS-CoV-2 strains, while also exhibiting significant differences with SARS and MERS (Figure #5.). We performed a similar alignment analysis of the HA gene of Influenza A and Influenza B to identify highly specific target sequences for these viruses (Figure #6 and 7 respectively.). All sequences were obtained from NCBI GenBank and compiled through BioEdit. Both DNA and RNA synthetic viral targets were synthesized via Integrated DNA Technologies (IDT). To target these viruses, we designed padlock probe (PLP) sequences by adding half of the reverse complement of the target sequence to each side of a standard and tested PLP backbone. For ligation to occur, a phosphate group modification was added to the 5’ end of the probe. A forward primer complementary to a region of the padlock probe backbone and a reverse primer that has the same sequence as another region of the padlock probe backbone were designed. Both the forward and reverse overlap primers have two phosphorothioate modifications at the 3’ end to prevent the exonuclease activity of phi29 from cutting the two primers. The reverse overlap primer synthesizes DNA strands using the complementary DNA strands synthesized from the forward primer as a template. The designed PLPs for each virus and the two primers usable for all PLPs were all synthesized by IDT.

**Figure #1.**
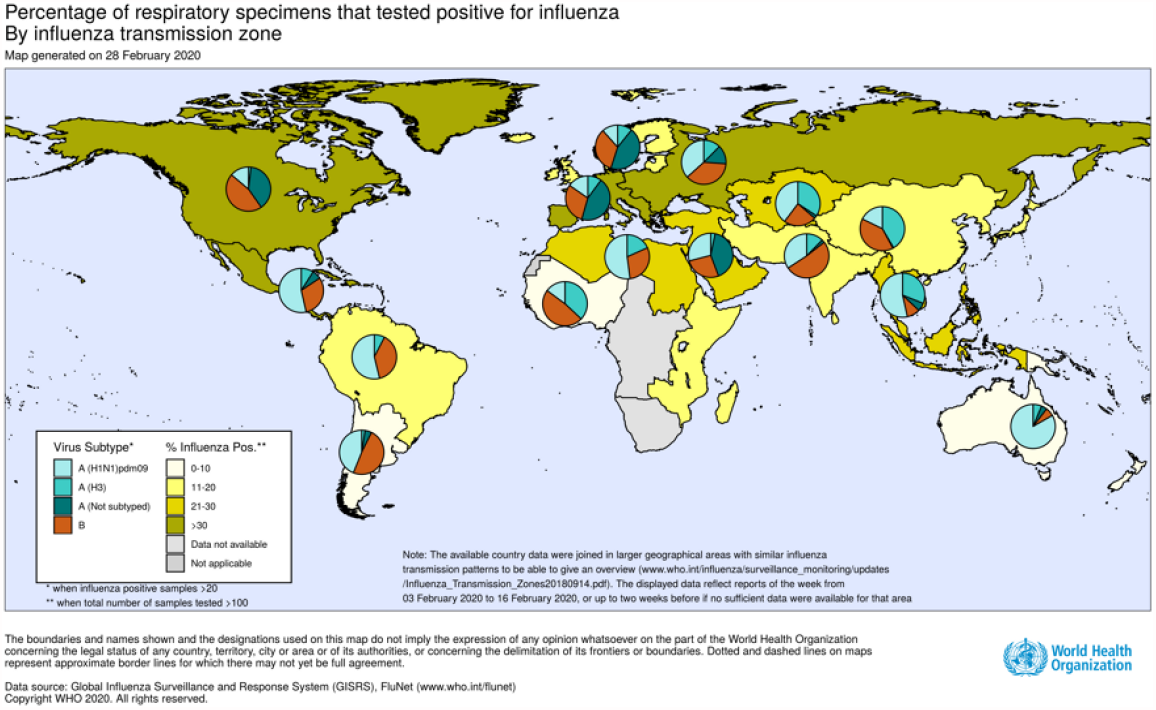
Influenza Cases Worldwide March 2020 (*WHO* | *Influenza Update - 362*, n.d.).

**Figure #2.**
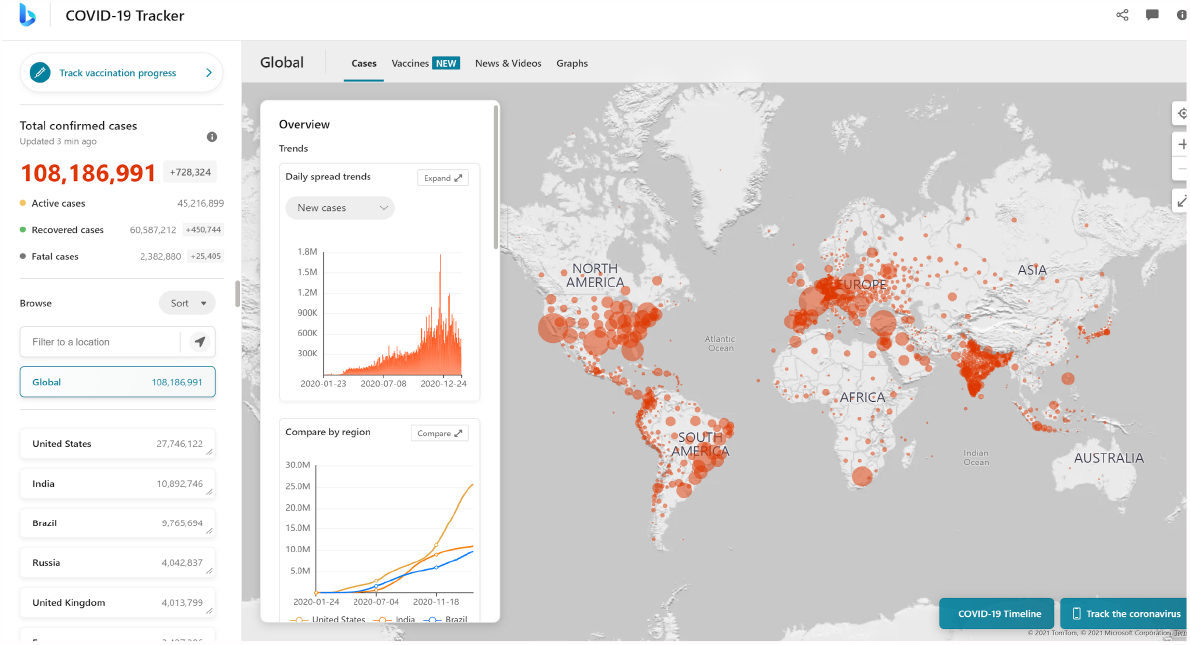
Worldwide SARS-CoV-2 Cases Up to February 13th 2021 (*Microsoft Bing COVID-19 Tracker*, n.d.).

**Figure #3.**
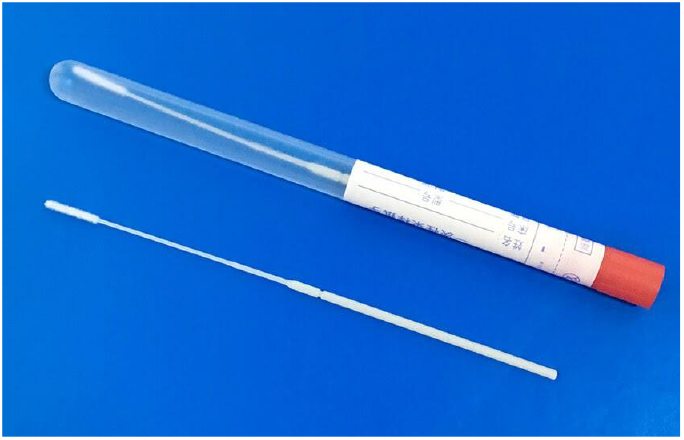
Nasopharyngeal Swab from Miraclean Technology (*Nasopharyngeal Sampling Swabs*, n.d.)

**Figure #4.**
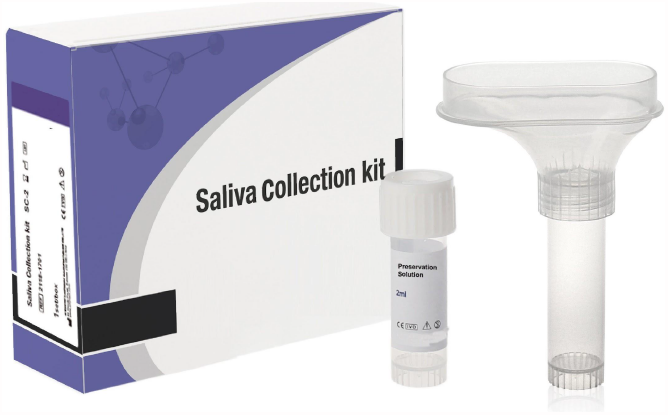
Saliva Collection Kit from Hangzhou Rollmed Co., Ltd. (*5ml Saliva Collection Tube Funnel Device Kit*, n.d.)

**Figure #5.**
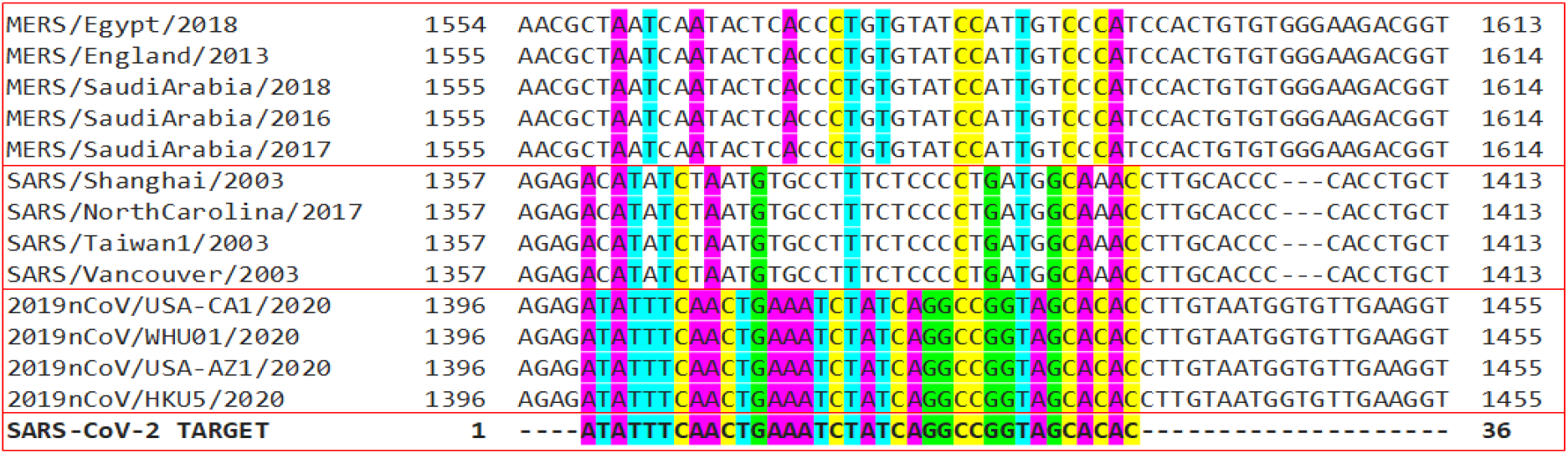
We chose the SARS-CoV-2 target sequence located in the Spike (S) gene as a target based on alignment data of various strains of SARS-CoV-2 (2019nCov), SARS, and MERS. We chose a 36 nucleotide sequence that is perfectly conserved with those of SARS-CoV2 while showing minimal matching with SARS and MERS.

**Figure #6.**
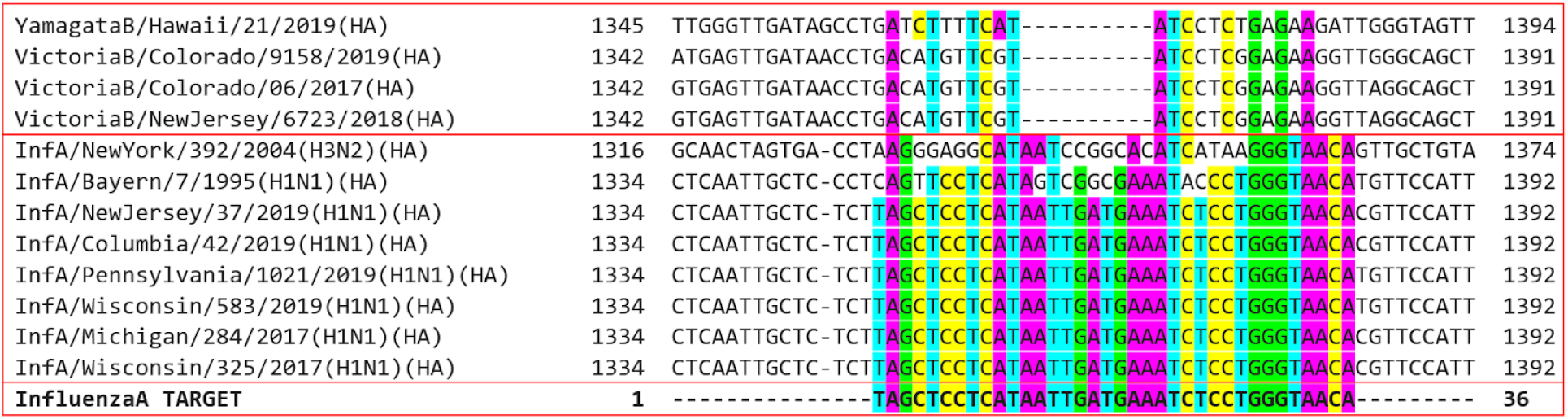
We chose the Influenza A target sequence located in the hemagglutinin gene as a target based on alignment data of various strains of Influenza A (H1N1pdm09) and Influenza B (Victoria Lineage). We chose a 36 nucleotide sequence that is perfectly conserved with those of H1N1 Influenza A while not matching those of Influenza B.

**Figure #7.**
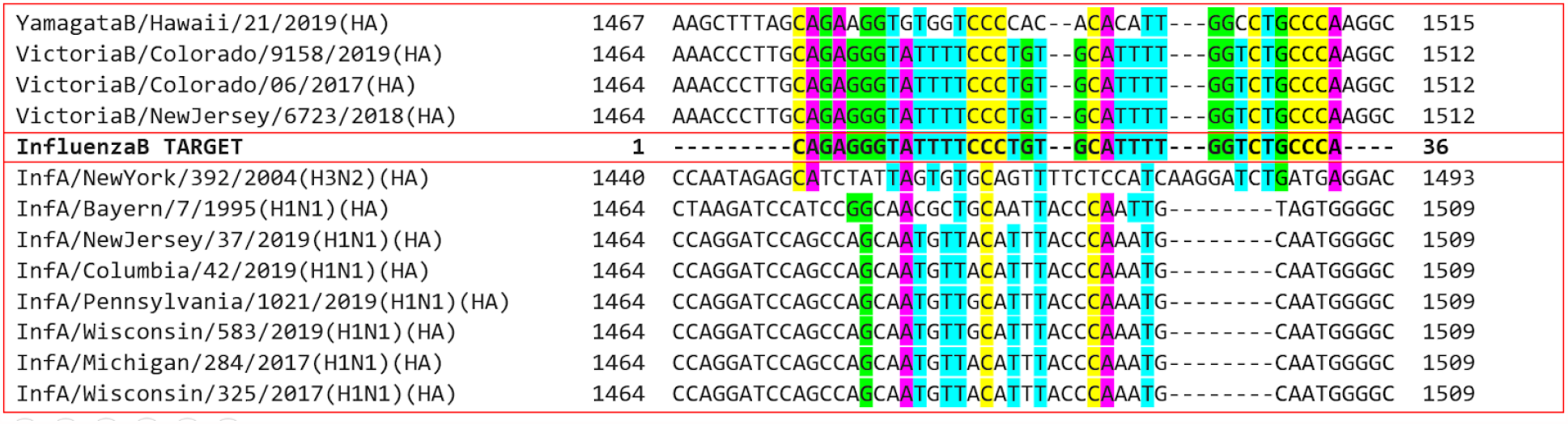
We chose the Influenza B target sequence located in the hemagglutinin gene as a target based on alignment data of various strains of Influenza A (H1N1pdm09) and Influenza B (Victoria Lineage). We chose a 36 nucleotide sequence that is perfectly conserved with those of H1N1 Influenza A while not matching those of Influenza B.

### Buffer-free reaction solutions

Due to this diagnostic test’s dependence on pH change, we created specific ligation and amplification reaction solutions with no buffers. These reaction solutions included the same reagents as the commercial reaction buffer from NEB but without Tris-HCl. Each 8X “no-Tris” ligation solution with 20 mM of MgCl_2_, 2 mM of ATP, 20 mM of DTT, 3.5 mM of NaOH, and deionized water (DI H_2_O). Similarly, each 2X “no-Tris” amplification solution with 20 mM of MgCl_2_, 20 mM of (NH_4_)_2_SO_4_, 8 mM of DTT, 2.8 mM of dNTP, 2 mM of NaOH, and DI H_2_O.

### Ligation and amplification reactions

The ligation and rolling circle amplification (RCA) reactions were run simultaneously in the same PCR tube at a total volume of 40 μL. For the ligation reaction, 5 μL of 8X “no-Tris” ligation solution was added to 1μL of a 2.5μM DNA or RNA target, 1μL of a 10μM padlock probe (PLP), 1μL of T4 ligase (for DNA targets) or splintR ligase (for RNA targets), 0.5μL of 10U Protector RNase Inhibitor (Sigma-Aldrich) (substituted with DI H_2_O for DNA targets), and 1.5μL of DI H_2_O. For the amplification reaction, 20μL of 2X “no-Tris” amplification solution, 2μL of 24μM forward primer (PLP-1), 2μL of 24μM reverse overlap primer (PLP-2), 4μL of 0.05% Phenol Red, and 2μL of 20U phi29 DNA polymerase were added to the 10μL ligation mixture. The PCR tube is placed at room temperature for at least 1.5 hours. For RCA testing of DNA targets, T4 ligase is utilized for the ligation of hybridized DNA padlock probe and DNA target junctions. For RCA testing of RNA targets, splintR ligase is utilized for the ligation of hybridized DNA padlock probe and RNA target junctions. Protector RNase Inhibitor was also utilized for RNA target testing to prevent RNA degradation.

### Confirmation of Nucleic Acid Amplification

Although there are no other known factors that could cause the same color change exhibited in our RCA reaction, we wanted to validate that our RCA test was functioning and changing colors due to DNA amplification. To confirm that DNA amplification was occuring, we performed RCA reactions where a positive target (where the RNA target matched the DNA padlock probe) was added and no target was added. We removed an aliquot of the reaction 30 min and 40 min into the reaction. Upon removal of each aliquot, we immediately heat shocked at 80°C to inactivate the enzymes and stop the RCA reaction. We then ran the samples on a 1% agarose gel to visualize any DNA.

### Sensitivity and Specificity Tests

To analyze the sensitivity of our assay, we selected the SARS CoV-2 RNA target and conducted serial dilutions down to nanomolar, picomolar, femtomolar, and attomolar concentrations of SARS-CoV-2 target to use in our RCA test. We utilized this method to determine the lowest viral RNA quantity that is still detectable by our RCA test.

To analyze the specificity of our assay, we determined whether differing sequences of synthetic viral targets that reflect different virus strains can be differentiated by our test. To find out, we used no target, a mismatched target, and a matched target for each of our tests as determined by our padlock probe (PLP). Each padlock probe has a specific sequence that is only fully complementary to its corresponding target, which should only instigate a reaction (Figure #8.).

**Figure #8.**
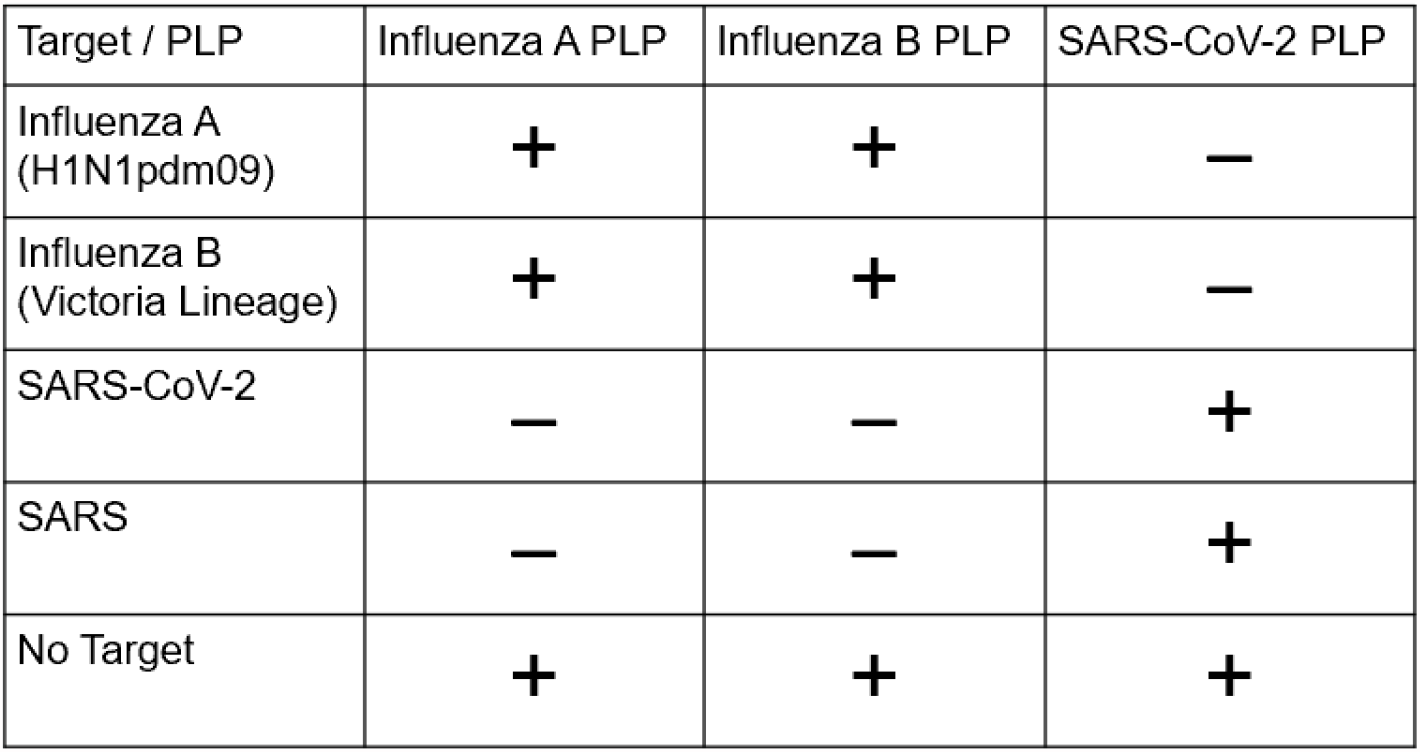
The table below shows the different synthetic RNA target and padlock probe combinations used in each reaction to test the specificity of our assay. (+) means the presence of the target while (-) means a lack thereof.

### Modeling Data Collection and Analysis

The volume of the reactions was insufficient for standard methods of spectrophotometry or pH measurement, and thus computer vision was used to quantify timelapses of the RCA reactions. Using the software we developed - VisualpH -pixels displaying the reactions were isolated in each frame of the time-lapse video, and values outside an empirically determined number of standard deviations from the mean were removed as outliers. The remaining pixels in each frame were averaged to plot mean hue against time. (Figure #9.).

**Figure #9.**
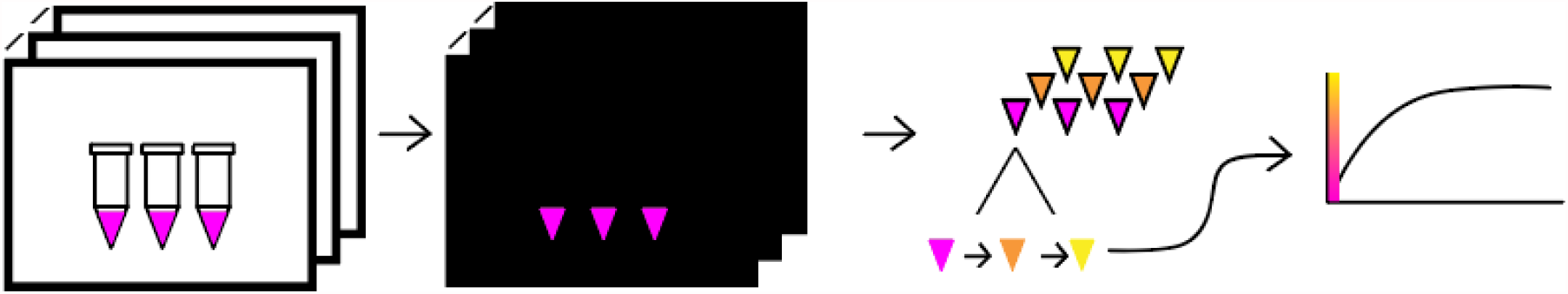
Masks are created by drawing polygons around the pixels representing the reaction. The masks are then applied to each frame, and the values of the isolated pixels are obtained and plotted against time.

**Figure #10.**
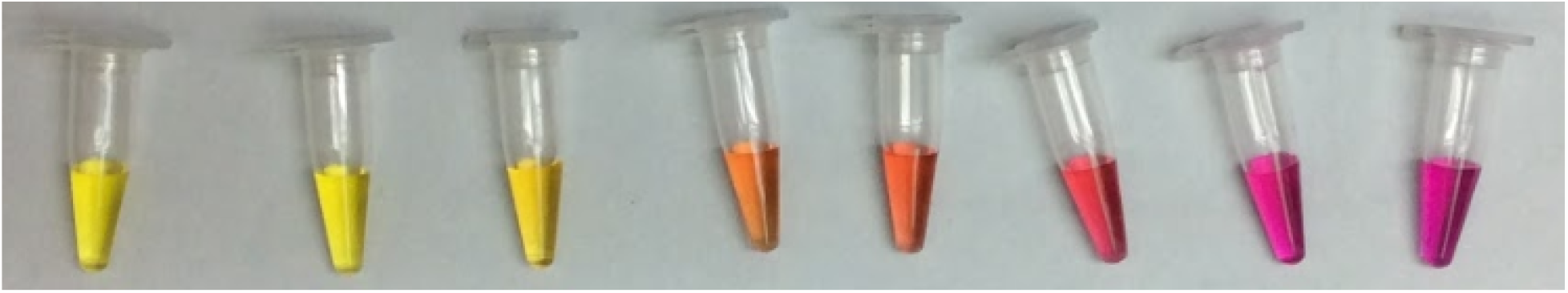
pH standards from left to right: 4.0, 5.0, 6.0, 7.0, 7.42, 8.0, 9.0, 10.0 showing the color change of phenol red across set pHs.

Using a set of pH standard solutions with the same phenol red indicator, both the pH and hue of the solutions were measured and plotted to determine a regression curve between the two values. VisualpH then converts the hue data into pH measurements to be plotted on a pH over time graph for all RCA reactions.

## Results and Discussion

### Confirmation of DNA Amplification

Our results show a significant amount of DNA remaining in the wells in the samples where we added a positive target (Figure #11.). In addition, these samples exhibited significant smearing of DNA across the entire lane of the gel. These results strongly suggest that DNA amplification occurred only in the samples where a positive target was added. Furthermore, the amplification resulted in highly branched and long nucleotide sequences, which prevented the samples from migrating through the gel. The smearing is likely due to a mixed population of varying nucleotide sequences. Lastly, we observed that the amount of DNA increased in the 40 minute time point compared to the 30 minute time point, which further supports our model that DNA amplification is occurring rapidly over time in our RCA reaction. During the amplification process, the incorporation of dNTPs into the sequence synthesized results in the formation of PPi and H^+^. As more and more H^+^ ions are made, the pH indicator changes from purple to yellow due to a drop in pH. Thus, these results demonstrate that the color change was most positively due to H^+^ concentration change that occurred via DNA amplification and not due to other factors unrelated to DNA synthesis.

**Figure #11.**
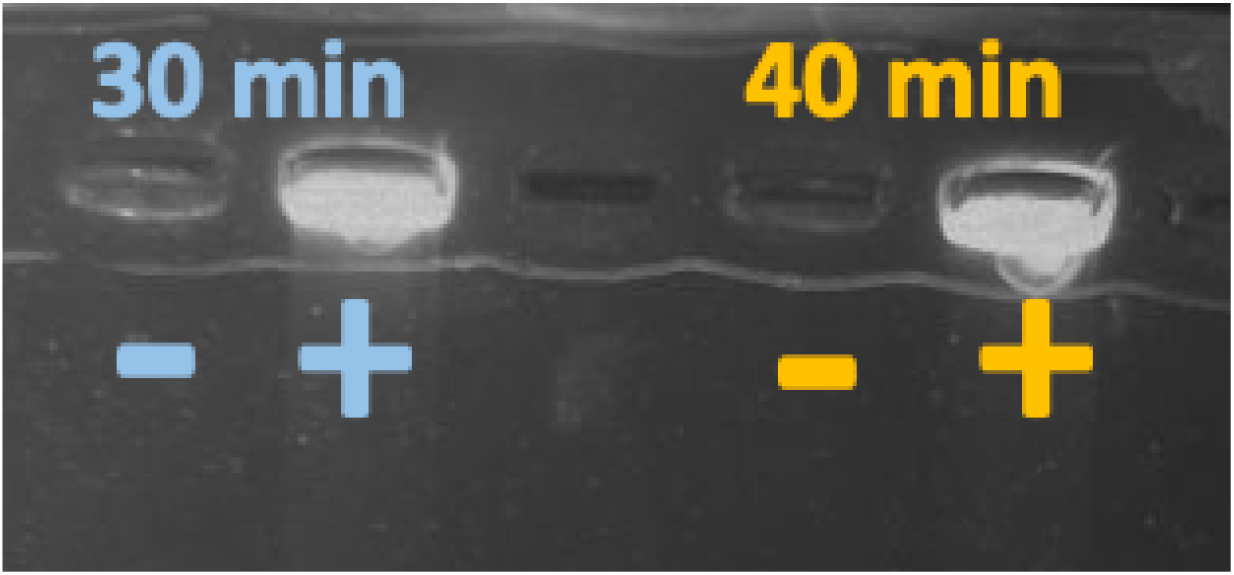
Comparison between viral load of SARS-CoV-2 after 30 minutes of amplification and 40 minutes of amplification. These samples were loaded with 6X concentrated DNA loading dye (geneaid) into 1% agarose gel.

**Figure #12.**
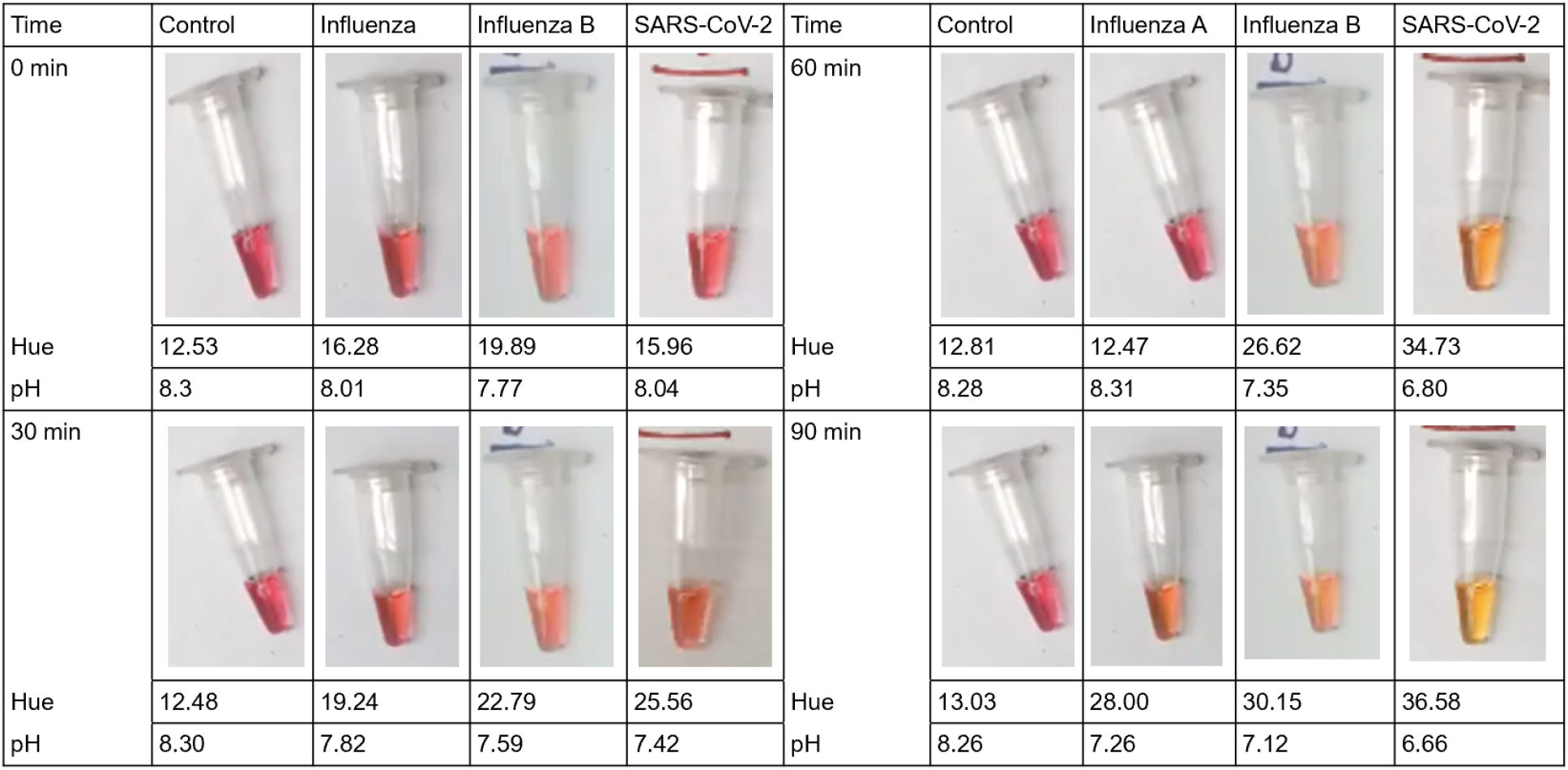
Image, hue, and pH of RCA reactions at time intervals 0, 30, 60, and 90 minute time intervals when detecting DNA synthetic viral targets.

**Figure #13.**
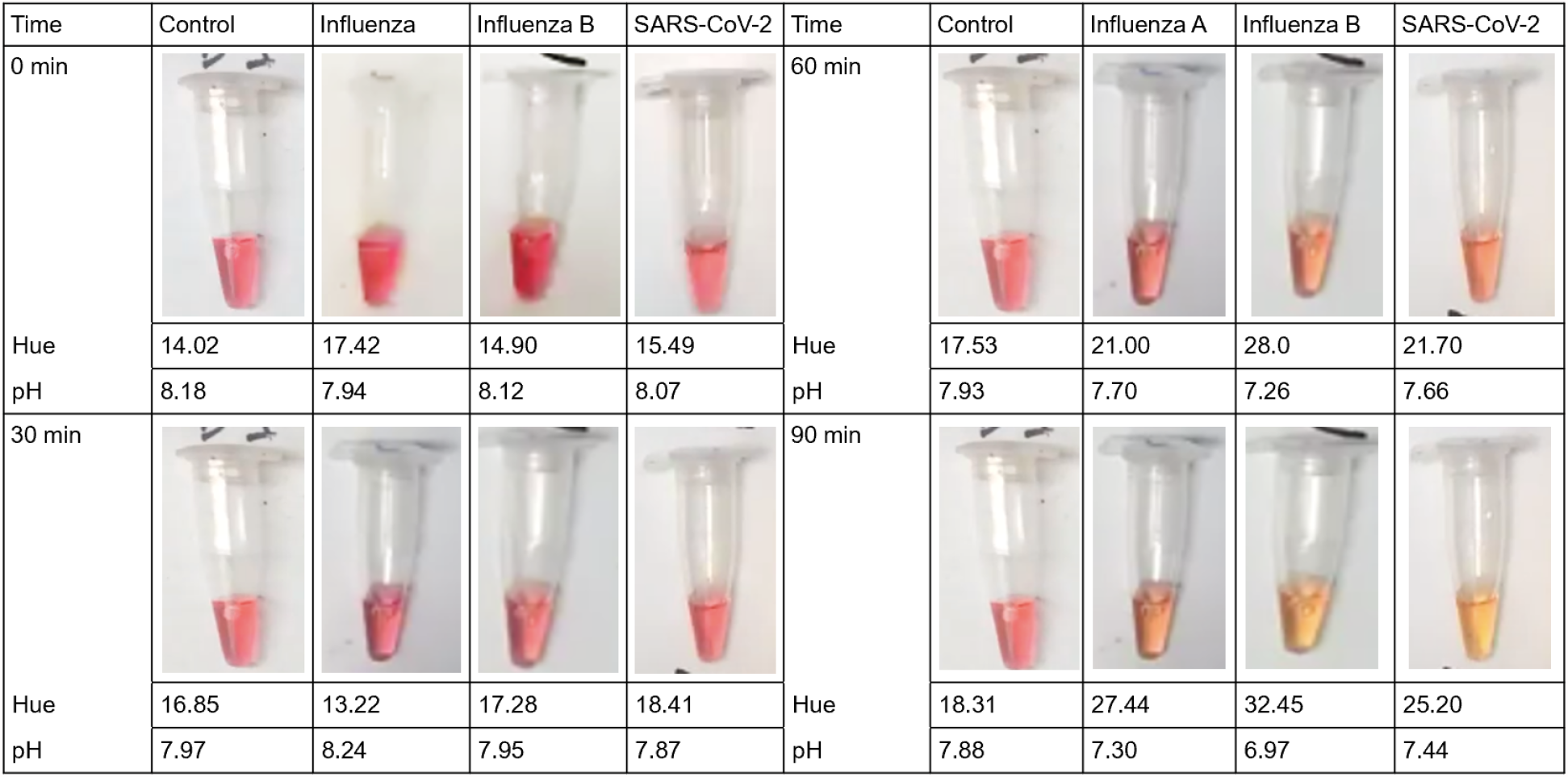
Image, hue, and pH of RCA reactions at time intervals 0, 30, 60, and 90 minute time intervals when detecting RNA synthetic viral targets.

### RCA Test using DNA and RNA Targets

Although our study aims to detect SARS-CoV-2, Influenza A (H1N1pdm09), and Influenza B (Victoria Lineage), which are all RNA viruses, testing DNA targets is also important as other notable DNA viruses including smallpox and papillomaviruses. This expands the possibility of our methodology to a wider array of viruses.

For this reason, the RCA reaction was run with both the RNA synthetic viral targets of Influenza A (INFA), Influenza B (INFB), and SARS-CoV-2, and the DNA form of the same targets as previously identified and selected. Using our software tool VisualpH, we were able to identify both the pH and hue associated with each reaction at any point in time and to create pH over time graphs.

Our results show that DNA testing for RCA typically yields faster color change results than that of RNA testing. The pH over time graph for Influenza A, Influenza B, and SARS-CoV-2, according to Figure # 14, 15, and 16 respectively, show that the DNA and the RNA test have similar trend lines. Although the pH of the DNA test (shown in orange) decreases faster than the RNA test (shown in blue) for some, our results show that both the DNA and RNA test decreases rapidly, which means our RCA test is functional with both synthetic DNA and RNA targets, specifically for our 3 chosen virus types.

**Figure #14.**
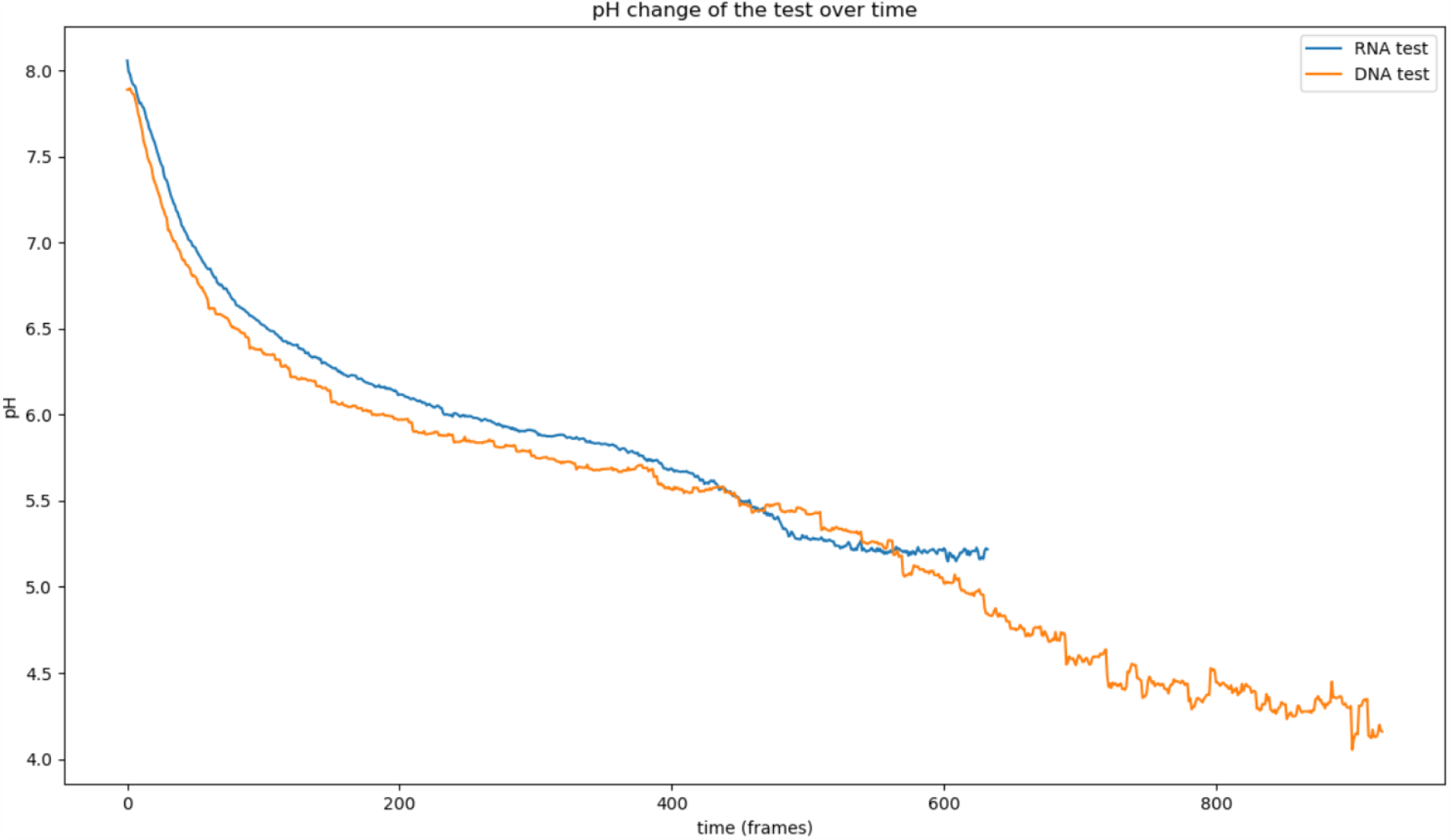
INFA test DNA versus RNA, a comparison between using the DNA and RNA test for Influenza A.

**Figure #15.**
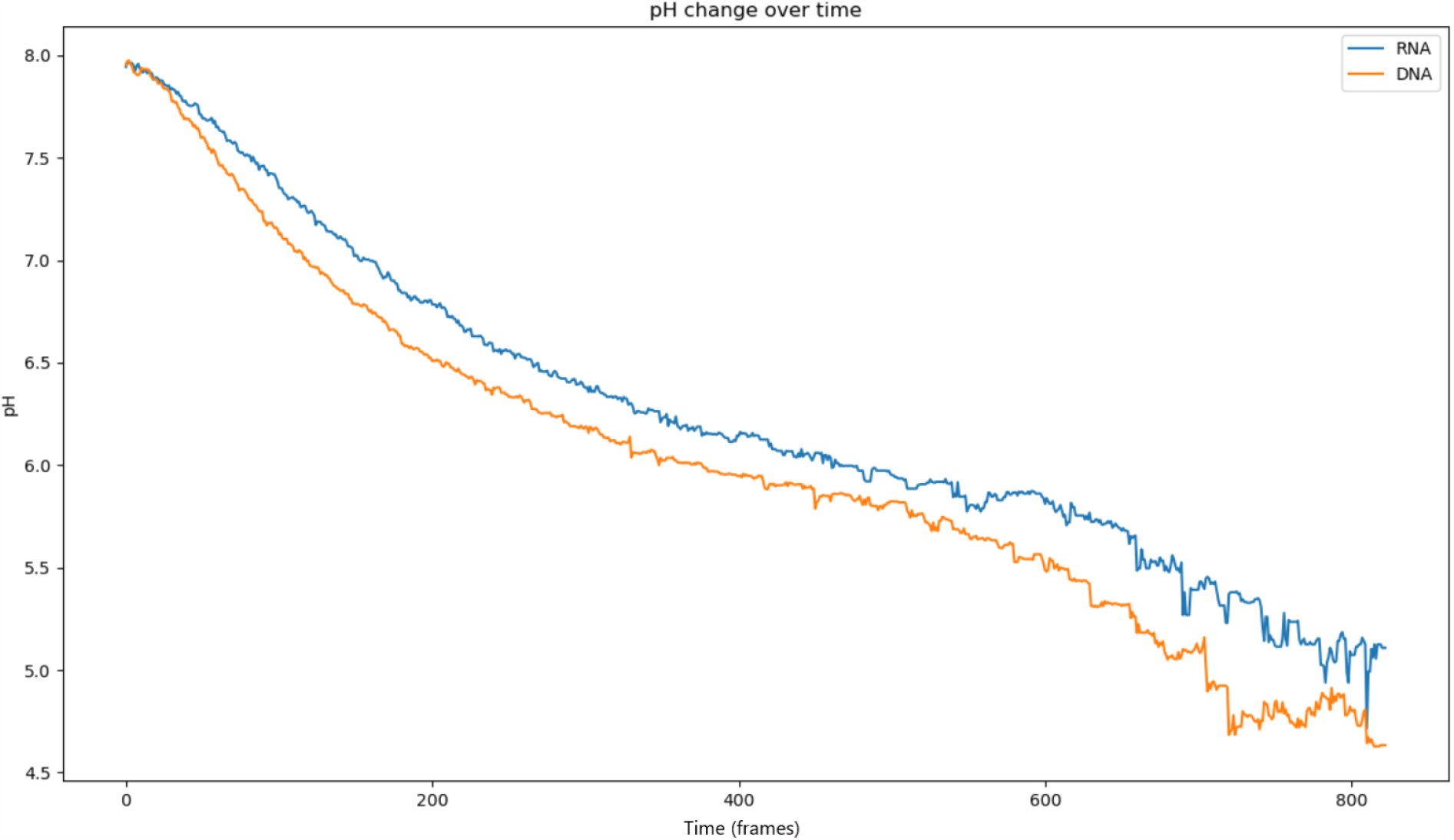
INFB test DNA versus RNA, a comparison between using the DNA and RNA test for Influenza B.

**Figure #16.**
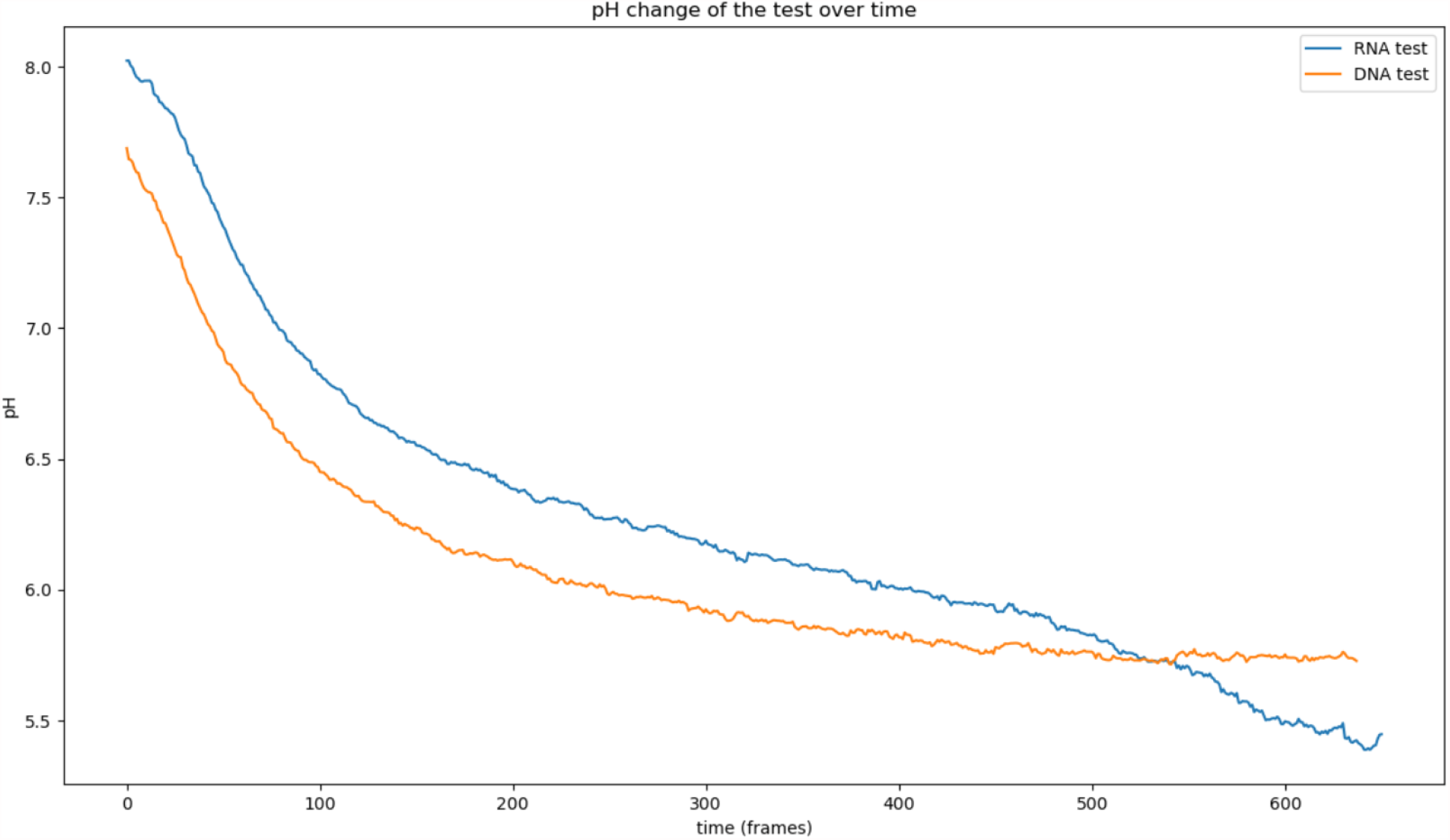
SARS-CoV-2 Test DNA vs RNA, a comparison between using the DNA and RNA Test for SARS-CoV-2.

**Figure #17.**
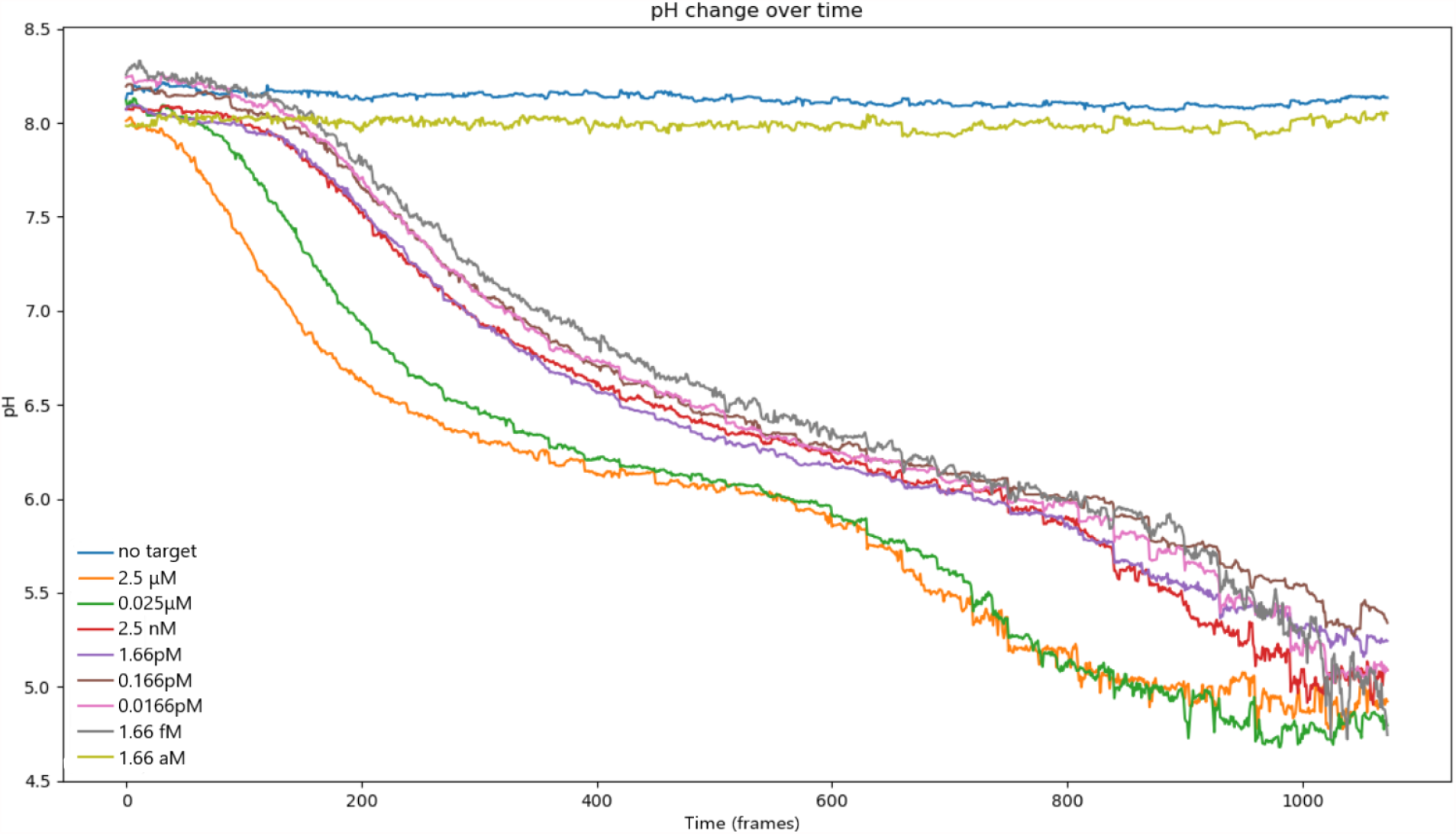
Sensitivity Test on SARS-CoV-2 target. Testing the sensitivity of our RCA test using different concentrations of SARS-CoV-2 targets in a 40μL RCA reaction.

**Figure #18.**
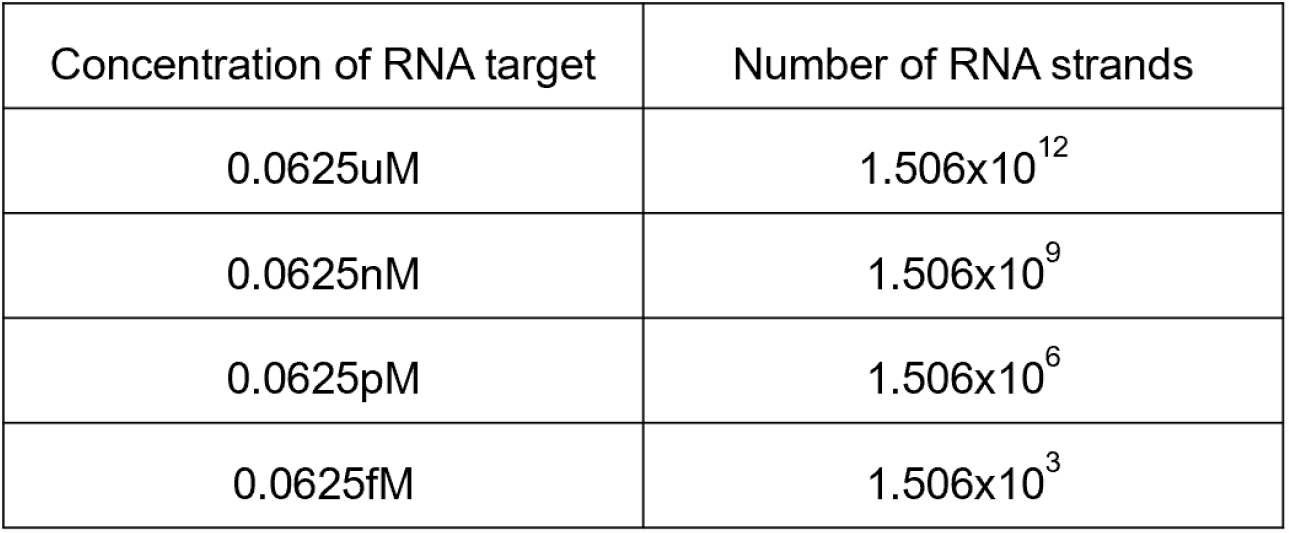
Conversion of RNA Concentration to Number of RNA strands.

**Figure #19.**
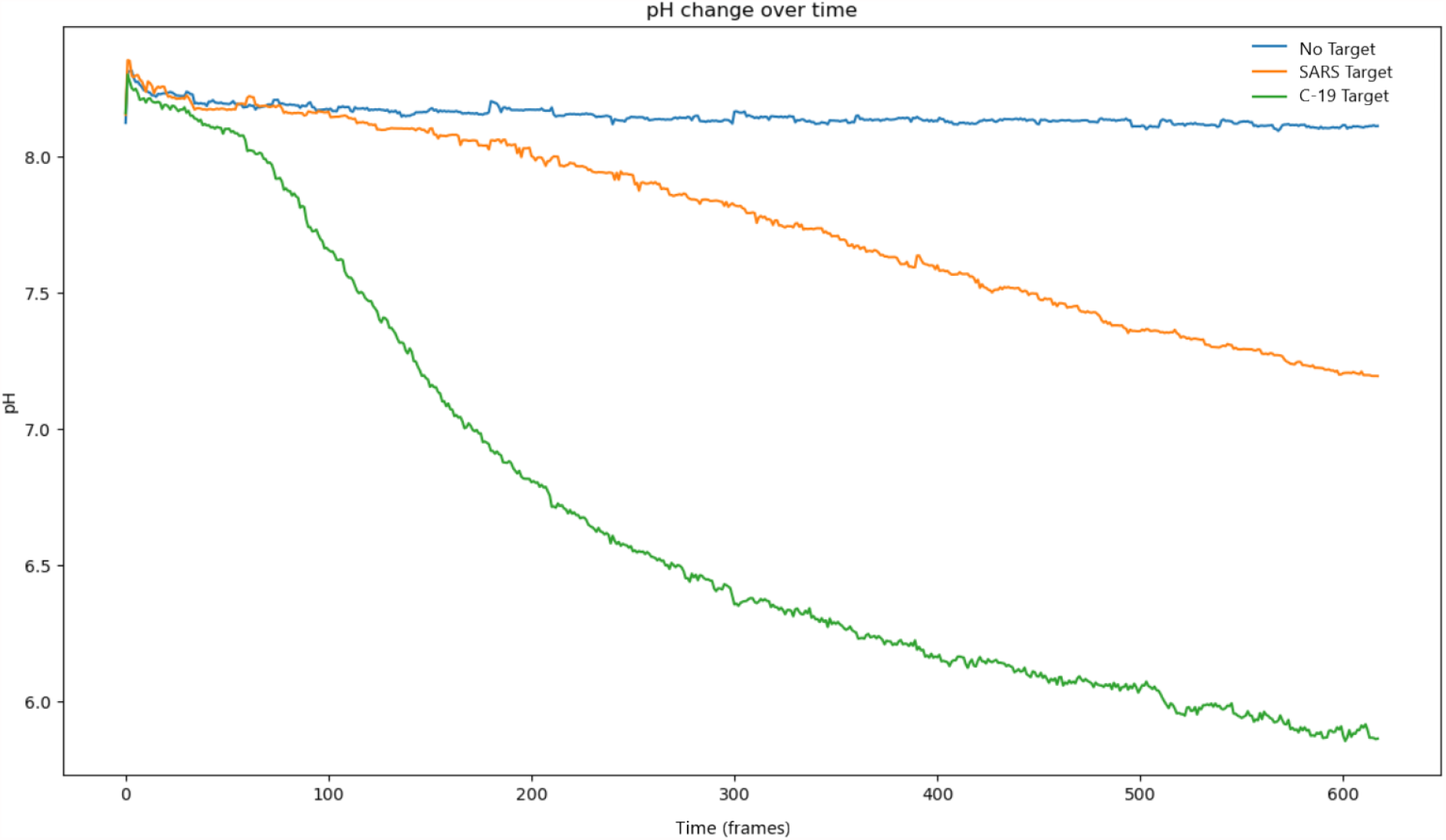
SARS-CoV-2 specificity test using the SARS-CoV-2 padlock probe against SARS, SARS-CoV-2, and no target in RCA reactions.

**Figure #20.**
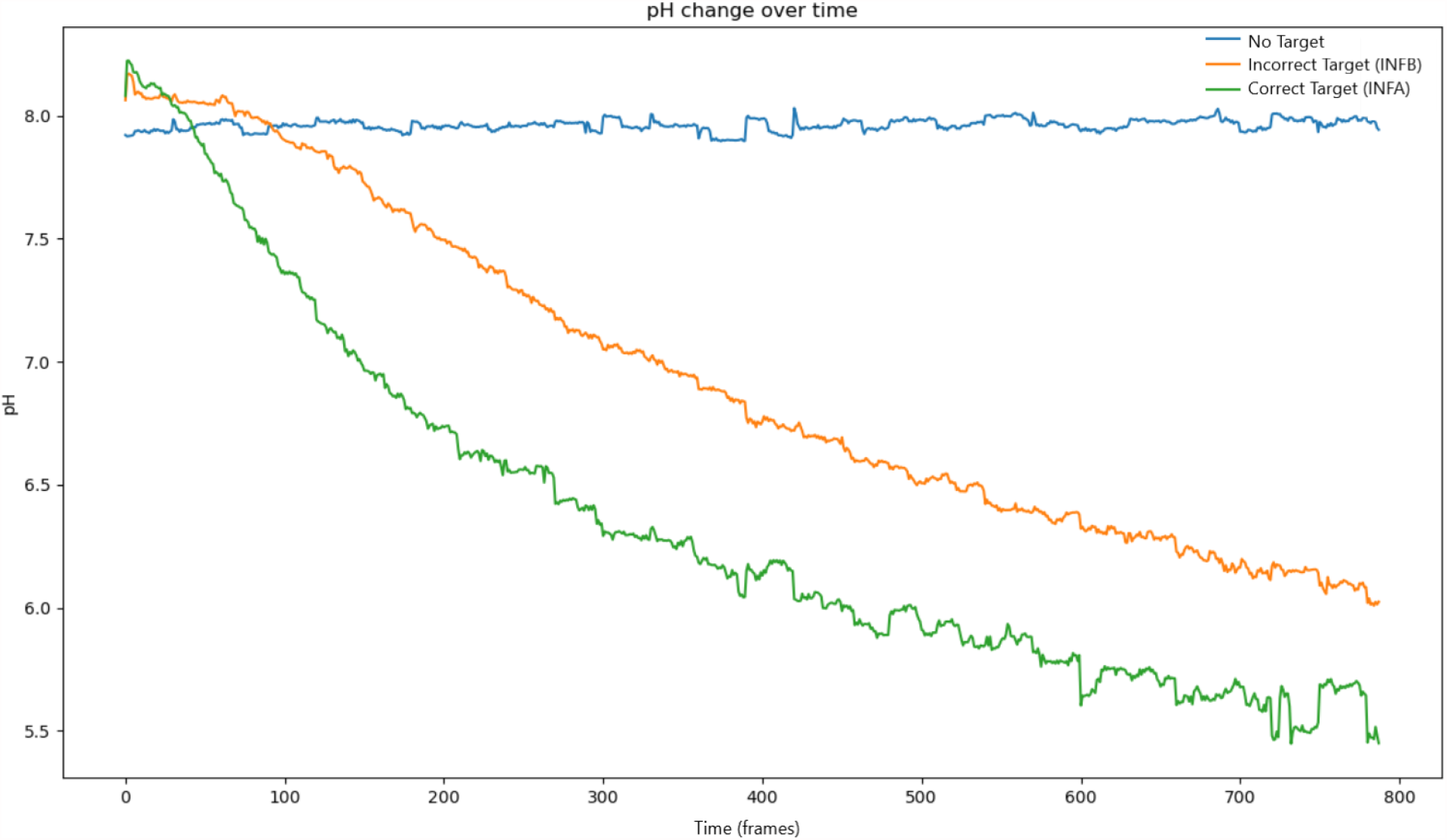
INFA specificity test using the INFA padlock probe against INFB, INFA, and no target in RCA reactions.

**Figure #21.**
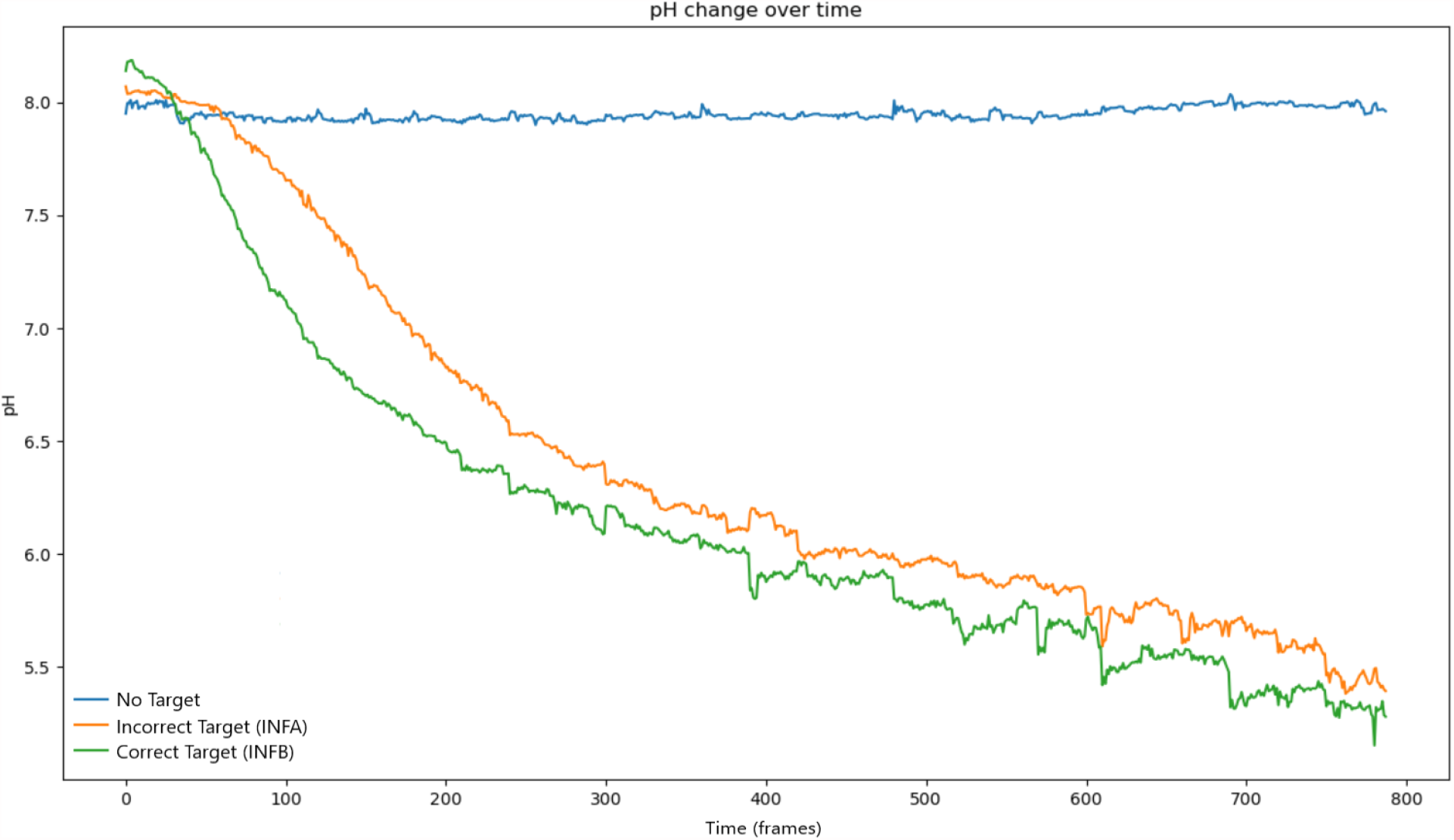
INFB specificity test using the INFB padlock probe against INFA, INFB, and no target in RCA reactions.

### Sensitivity Tests

Our results demonstrate that a color change occurs in all concentrations until the attomolar concentration is reached. Though some of the lower concentrations relied on a longer reaction time, it shows that our assay is highly sensitive in detecting synthetic viral RNA strands (Figure #17.). Specifically, we can still detect up to 0.0415pM of SARS-CoV-2 target or around 1000 synthetic strands while differentiating from a negative test. The below table indicates some of the concentrations of synthetic RNA targets in terms of RNA strands.

### Specificity Tests

Our results show that when the SARS-CoV-2 padlock probe is used, the pH of the SARS-CoV-2 target solution has the greatest change. As expected, the solution with no target has little to no pH change. The SARS target has a slight pH change due to its similarities with the SARS-CoV-2 nucleotide sequence, but the color difference is still significant enough to distinguish between the two.

Our results highlight the fact that our padlock probe could detect the difference between the different targets. Although the color and pH of both targets change, the rate of change for the correct target was much faster than the incorrect target.

Our results highlight the fact that our padlock probe could detect the difference between the different targets. Although the color and pH of both targets change, the rate of change for the correct target was much faster than the incorrect target.

Despite similar nucleotide sequences, a mismatched viral sequence will result in a slower color change than that of when the virus sequence actually tested is present. In our SARS-CoV-2 test, the presence of SARS-CoV-2 target sequence resulted in a notably faster reaction time than in the presence of a SARS target sequence. Similar results were found for the Influenza tests, using the target combinations listed above. The notable time difference in reaction times allows us to differentiate different viruses in the same type, whether that is the influenza virus or the coronavirus.

## Further Directions

### Lentivirus and Live Virus Testing

There are various limitations in our study that should be addressed. One of them being the feasibility of this methodology for an actual diagnostic test. For our RCA tests, we utilized 36bp synthetic DNA and RNA targets as outlined in the *Synthetic Viral Targets, Probes and Primers* section. As the entire gene sequence from the targeted region usually contains thousands of base pairs and the entire genome of the targeted virus contains tens of thousands of base pairs, it would theoretically be much more difficult for our RCA platform to detect the targeted 36bp. As such, the sensitivity and specificity of our RCA tests in a real diagnostic setting is likely to be much lower than our shown results. The degree of this potential impact could be determined by conducting RCA tests using the isolated nucleic acid of lentiviruses that model our targeted virus. Testing with the isolated nucleic acid of the actual live virus in P2 labs could further solidify the findings. Doing these additional tests could help evaluate the possible further modifications necessary to create a RCA platform.

### Point of Care Sample Preparation

This methodology only presents the DNA amplification and detection portion of a diagnostic test. Since this is a nucleic acid test, we would likely require sample preparation procedures to isolate either the DNA or RNA of the virus. The most utilized lab technique includes the use of DNA or RNA purification kits. However, these are not as viable in a non-lab and point of care setting. A few novel techniques are currently proposed to solve this issue. One method involves a simple syringe and membrane filter system that could purify nucleic acids from pathogens in as little as 20 minutes (Zhao et al., 2019). Another method is a hardware designed by Akonni Biosystems called TruTip Nucleic Acid Purification which conducts DNA or RNA extractions and delivers inhibitor-free, PCR-ready products in as quickly as 4 minutes (Thakore et al., 2018).

## References

Alhassan, Andy, Zhiru Li, Catherine B. Poole, and Clotilde K.S. Carlow. “Expanding the MDx Toolbox for Filarial Diagnosis and Surveillance.” Trends in Parasitology 31, no. 8 (August 2015): 391–400. https://doi.org/10.1016/j.pt.2015.04.006.

American College of Cardiology. “False-Negative Rate of RT-PCR SARS-CoV-2 Tests.” Accessed June 5, 2020. %3a%2f%2f www.acc.org%2flatest-in-cardiology%2fjournal-scans%2f2020%2f05

Corman, Victor M, Olfert Landt, Marco Kaiser, Richard Molenkamp, Adam Meijer, Daniel KW Chu, Tobias Bleicker, et al. “Detection of 2019 Novel Coronavirus (2019-NCoV) by Real-Time RT-PCR.” Eurosurveillance 25, no. 3 (January 23, 2020). https://doi.org/10.2807/1560-7917.ES.2020.25.3.2000045.

Deng, Ruijie, Kaixiang Zhang, Yupeng Sun, Xiaojun Ren, and Jinghong Li. “Highly Specific Imaging of MRNA in Single Cells by Target RNA-Initiated Rolling Circle Amplification.” Chemical Science 8, no. 5 (2017): 3668–75. https://doi.org/10.1039/C7SC00292K.

Drosten, Christian, Stephan Günther, Wolfgang Preiser, Sylvie van der Werf, Hans-Reinhard Brodt, Stephan Becker, Holger Rabenau, et al. “Identification of a Novel Coronavirus in Patients with Severe Acute Respiratory Syndrome.” New England Journal of Medicine 348, no. 20 (May 15, 2003): 1967–76. https://doi.org/10.1056/NEJMoa030747.

DziŐbowska, Karolina, Elżbieta Czaczyk, and Dawid Nidzworski. “Detection Methods of Human and Animal Influenza Virus—Current Trends.” Biosensors 8, no. 4 (October 18, 2018): 94. https://doi.org/10.3390/bios8040094.

Fink, C. G. “Molecular Methods for Virus Detection.” Molecular Pathology 50, no. 2 (April 1, 1997): 111–111. https://doi.org/10.1136/mp.50.2.111-a.

Franzén, Mikael. “Covid-19 Rapid Test - BLUE PAPER.” Preprint. Open Science Framework, March 10, 2020. https://doi.org/10.31219/osf.io/jpukc.

Ge, Yiyue, Bin Wu, Xian Qi, Kangchen Zhao, Xiling Guo, Yefei Zhu, Yuhua Qi, et al. “Rapid and Sensitive Detection of Novel Avian-Origin Influenza A (H7N9) Virus by Reverse Transcription Loop-Mediated Isothermal Amplification Combined with a Lateral-Flow Device.” Edited by Xue-jie Yu. PLoS ONE 8, no. 8 (August 1, 2013): e69941. https://doi.org/10.1371/journal.pone.0069941.

Ge, Yiyue, Qiang Zhou, Kangchen Zhao, Ying Chi, Bin Liu, Xiaoyan Min, Zhiyang Shi, Bingjie Zou, and Lunbiao Cui. “Detection of Influenza Viruses by Coupling Multiplex Reverse-Transcription Loop-Mediated Isothermal Amplification with Cascade Invasive Reaction Using Nanoparticles as a Sensor.” International Journal of Nanomedicine Volume 12 (April 2017): 2645–56. https://doi.org/10.2147/IJN.S132670.

Gu, Lide, Wanli Yan, L. Liu, Shujun Wang, Xu Zhang, and Mingsheng Lyu. “Research Progress on Rolling Circle Amplification (RCA)-Based Biomedical Sensing.” Pharmaceuticals 11, no. 2 (April 21, 2018): 35. https://doi.org/10.3390/ph11020035.

Gyarmati, P., T. Conze, S. Zohari, N. LeBlanc, M. Nilsson, U. Landegren, J. Baner, and S. Belak. “Simultaneous Genotyping of All Hemagglutinin and Neuraminidase Subtypes of Avian Influenza Viruses by Use of Padlock Probes.” Journal of Clinical Microbiology 46, no. 5 (May 1, 2008): 1747–51. https://doi.org/10.1128/JCM.02292-07.

Hamidi, Seyed Vahid, and Hedayatollah Ghourchian. “Colorimetric Monitoring of Rolling Circle Amplification for Detection of H5N1 Influenza Virus Using Metal Indicator.” Biosensors and Bioelectronics 72 (October 2015): 121–26. https://doi.org/10.1016/j.bios.2015.04.078.

Hamidi, Seyed Vahid, and Jonathan Perreault. “Simple Rolling Circle Amplification Colorimetric Assay Based on PH for Target DNA Detection.” Talanta 201 (August 2019): 419–25. https://doi.org/10.1016/j.talanta.2019.04.003.

Hartman, Mark R., Roanna C. H. Ruiz, Shogo Hamada, Chuanying Xu, Kenneth G. Yancey, Yan Yu, Wei Han, and Dan Luo. “Point-of-Care Nucleic Acid Detection Using Nanotechnology.” Nanoscale 5, no. 21 (2013): 10141. https://doi.org/10.1039/c3nr04015a.

Khan, Muhammad Imran, Koushik Mukherjee, Rizwan Shoukat, and Huang Dong. “A Review on PH Sensitive Materials for Sensors and Detection Methods.” Microsystem Technologies 23, no. 10 (October 2017): 4391–4404. https://doi.org/10.1007/s00542-017-3495-5.

Larsson, Chatarina, Ida Grundberg, Ola Söderberg, and Mats Nilsson. “In Situ Detection and Genotyping of Individual MRNA Molecules.” Nature Methods 7, no. 5 (May 2010): 395–97. https://doi.org/10.1038/nmeth.1448.

Larsson, Chatarina, Jørn Koch, Anders Nygren, George Janssen, Anton K Raap, Ulf Landegren, and Mats Nilsson. “In Situ Genotyping Individual DNA Molecules by Target-Primed Rolling-Circle Amplification of Padlock Probes.” Nature Methods 1, no. 3 (December 2004): 227–32. https://doi.org/10.1038/nmeth723.

Lippi, Giuseppe, Camilla Mattiuzzi, Chiara Bovo, and Mario Plebani. “Current Laboratory Diagnostics of Coronavirus Disease 2019 (COVID-19).” Acta Bio Medica Atenei Parmensis 91, no. 2 (May 11, 2020): 137–45. https://doi.org/10.23750/abm.v91i2.9548.

Lohman, Gregory J. S., Yinhua Zhang, Alexander M. Zhelkovsky, Eric J. Cantor, and Thomas C. Evans. “Efficient DNA Ligation in DNA–RNA Hybrid Helices by Chlorella Virus DNA Ligase.” Nucleic Acids Research 42, no. 3 (February 2014): 1831–44. https://doi.org/10.1093/nar/gkt1032.

“Rapid Influenza Diagnostic Tests | CDC,” November 12, 2019. https://www.cdc.gov/flu/professionals/diagnosis/clinician_guidance_ridt.htm.

Roberts, J., K Bebenek, and T. Kunkel. “The Accuracy of Reverse Transcriptase from HIV-1.” Science 242, no. 4882 (November 25, 1988): 1171–73. https://doi.org/10.1126/science.2460925.

Rota, P. A. “Characterization of a Novel Coronavirus Associated with Severe Acute Respiratory Syndrome.” Science 300, no. 5624 (May 30, 2003): 1394–99. https://doi.org/10.1126/science.1085952.

Schott, Juliane W, Michael Morgan, Melanie Galla, and Axel Schambach. “Viral and Synthetic RNA Vector Technologies and Applications.” Molecular Therapy 24, no. 9 (September 2016): 1513–27. https://doi.org/10.1038/mt.2016.143.

Sexton, Tom, Sreenivasulu Kurukuti, Jennifer A Mitchell, David Umlauf, Takashi Nagano, and Peter Fraser. “Sensitive Detection of Chromatin Coassociations Using Enhanced Chromosome Conformation Capture on Chip.” Nature Protocols 7, no. 7 (July 2012): 1335–50. https://doi.org/10.1038/nprot.2012.071.

Thakore, Nitu, Ryan Norville, Molly Franke, Roger Calderon, Leonid Lecca, Michael Villanueva, Megan B. Murray, Christopher G. Cooney, Darrell P. Chandler, and Rebecca C. Holmberg. “Automated TruTip Nucleic Acid Extraction and Purification from Raw Sputum.” Edited by Sylvia Bruisten. PLOS ONE 13, no. 7 (July 5, 2018): e0199869. https://doi.org/10.1371/journal.pone.0199869.

Thuronyi, Benjamin W., Luke W. Koblan, Jonathan M. Levy, Wei-Hsi Yeh, Christine Zheng, Gregory A. Newby, Christopher Wilson, et al. “Continuous Evolution of Base Editors with Expanded Target Compatibility and Improved Activity.” Nature Biotechnology 37, no. 9 (September 2019): 1070–79. https://doi.org/10.1038/s41587-019-0193-0.

Vashist, Sandeep Kumar. “In Vitro Diagnostic Assays for COVID-19: Recent Advances and Emerging Trends.” Diagnostics 10, no. 4 (April 5, 2020): 202. https://doi.org/10.3390/diagnostics10040202.

Vega, M. de, J. M. Lazaro, M. Mencia, L. Blanco, and M. Salas. “Improvement of 29 DNA Polymerase Amplification Performance by Fusion of DNA Binding Motifs.” Proceedings of the National Academy of Sciences 107, no. 38 (September 21, 2010): 16506–11. https://doi.org/10.1073/pnas.1011428107.

Zedalis, Julianne, John Eggebrecht, and OpenStax College. Biology for AP® Courses, 2017. https://openstax.org/details/books/biology-ap-courses.

